# Pervasive RNA splicing in a plant DNA virus

**DOI:** 10.1101/2025.10.01.679800

**Authors:** Delphine M. Pott, Man Gao, Chaonan Shi, Liping Wang, Laura Medina-Puche, Zara Yagci, Sarah A. Acatay, Laura Hohenstein, Jun Zhang, Lucía Gonzalo, Catharina Merchante, Lianfeng Gu, Rosa Lozano-Durán

## Abstract

RNA splicing is considered an oddity in plant viruses; here, unexpectedly, we find it to be prevalent in the geminivirus tomato yellow leaf curl virus. Transcriptome analysis revealed eight splicing events generating novel protein isoforms, six of which were validated and two shown to depend on the plant spliceosome. Splicing impairment reduced viral accumulation and symptom severity, demonstrating that splicing contributes to infectivity, potentially by expanding the viral proteome, and suggesting that plant virus transcriptomes need to be reassessed.

## MAIN TEXT

Due to their limited coding space, viruses have evolved strategies to maximize coding capacity and efficiently carry out their functions. Splicing of viral transcripts, for example, increases the diversity of the viral proteome in mammalian viruses replicating in the nucleus (Li et al., 2023; Meyer, 2016). In plant viruses, however, splicing is assumed to be extremely rare, with only a few examples in members of the family *Geminiviridae* (genera *Mastrevirus*, *Becurtovirus*, and *Grablovirus*), with one potential splicing event described in each case, as well as *Caulimoviridae* (Shivaprasad et al., 2005; Wang & Lozano-Durán, 2023). In the absence of a systematic investigation, nevertheless, the *de facto* prevalence of splicing in viral transcripts in plants remains to be determined. By revisiting the prevailing view that plant virus transcripts are not generally spliced, our study provides evidence that alternative splicing may be more prevalent than previously appreciated.

To assess the incidence of splicing events affecting viral transcripts in plants, we re-analyzed RNA sequencing data of *Nicotiana benthamiana* leaves infected by the geminivirus tomato yellow leaf curl virus (TYLCV) (Wang et al., 2022). TYLCV belongs to the largest genus in the *Geminiviridae* family, *Begomovirus*, whose members have not been described to utilize splicing to date. Strikingly, we identified eight putative splicing events affecting viral transcripts produced from both DNA strands, of which six contained the most abundant canonical splice sites (GT-AG) at the intron boundaries, and two the second most common splice sites and hallmark for alternative splicing (GC-AG) (Figure 1A; Supplementary table 1). In most cases, these events are predicted to generate novel viral protein isoforms, by removing part of the protein sequence or creating frameshifts (Supplementary table 1).

**Figure 1.**
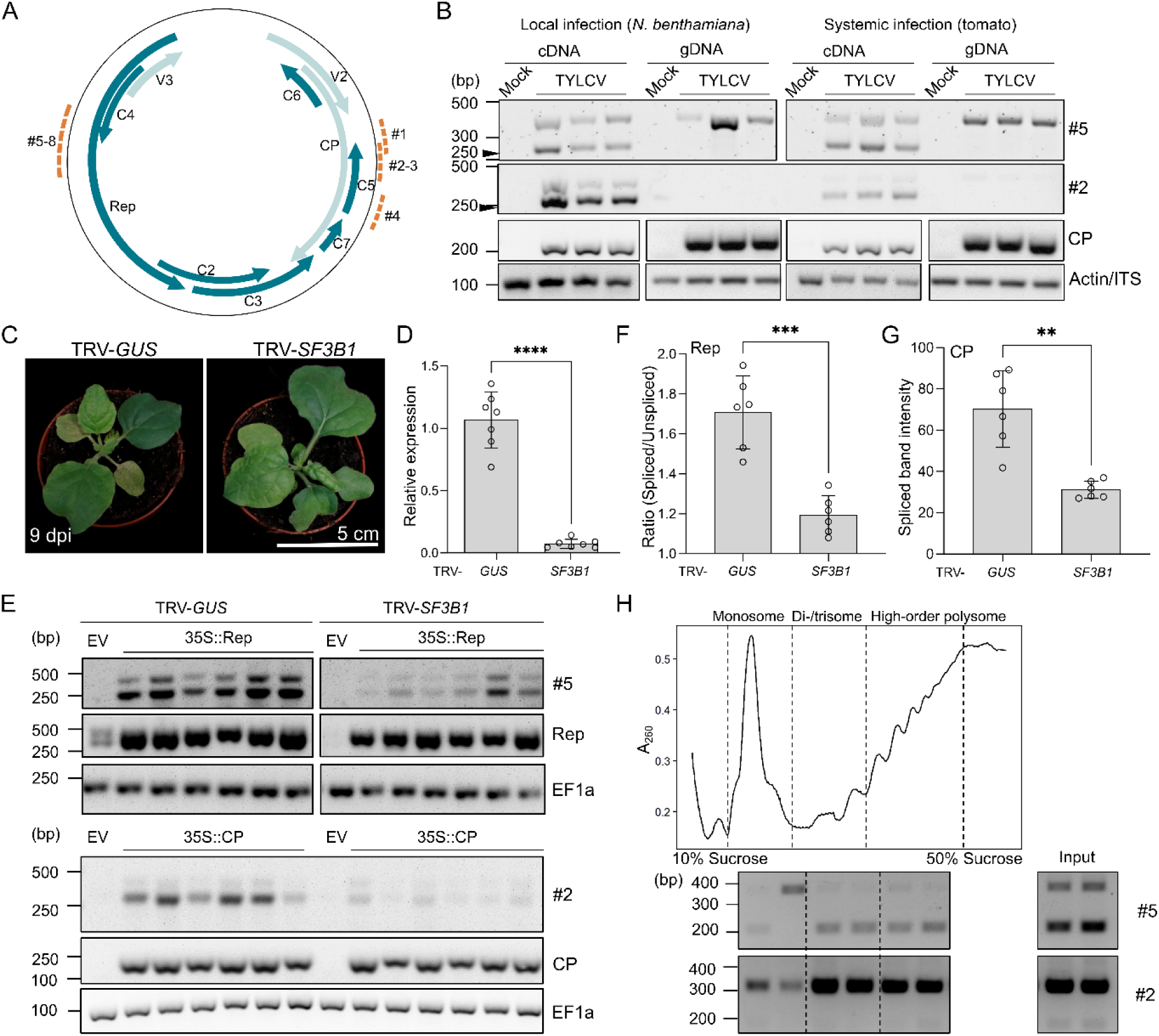
TYLCV transcripts undergo splicing *in planta*. (A) Representation of TYLCV genome; dashed orange lines: splicing events, light and dark green arrows: genes in the viral and complementary strand, respectively. (B) (RT-)PCR of viral cDNA and gDNA showing Rep/CP spliced transcripts (splicing events #5 and #2, respectively, indicated by arrowheads). (C) Phenotype of *SF3B1-*silenced plants (TRV-*SF3B1*) at 9 days post-inoculation (dpi). (D) *SF3B1* relative expression in TRV-*GUS-* and TRV-*SF3B1-*infected plants. (E) Splicing of CP/Rep (transiently expressed under the control of the constitutive cauliflower mosaic virus 35S promoter) in TRV-*GUS-* and TRV-*SF3B1-*infected plants. (F) Quantification of the ratio of spliced-to-unspliced transcripts (for Rep, as the primers amplify both spliced and unspliced transcripts) and (G) spliced band intensity (for CP; only the spliced transcript is reliability detected after RT-PCR) in TRV-*GUS-* and TRV-*SF3B1-*infected plants. (H) Spliced transcripts detected by polysome profiling in TYLCV-infected tomato plants. RT-PCR to detect splicing events #5 and #2 (lower) were performed in the different ribosomal fractions (upper) and input of two independent replicates.

As a proof of concept, we selected two of these splicing events, #5 (affecting the Rep transcript, encoding the replication-associated protein – complementary strand) and #2 (affecting the CP transcript, encoding the capsid protein – viral strand), for validation. Through PCR, we detected the corresponding spliced transcripts from cDNA but not from DNA isolated from TYLCV-infected samples, indicating that this modification takes place at the level of the RNA and is not the result of genomic rearrangements (Figure 1B). The identity of the spliced transcripts was confirmed by Sanger sequencing and using primers either flanking the splicing events or designed at the “exon-intron” junction (Supplementary Figure 1A-B). In addition to splicing events #2 and #5, we could also confirm events #1, #3, #4 and #8 following a similar strategy (Supplementary Figure 1C). Silencing of the core spliceosome component SF3B1 (Butt et al., 2021; Sun, 2020) by virus-induced gene silencing (VIGS) (Figure 1C, D) significantly decreased events #2 and #5 (Figure 1E-G), consistent with the idea that these variants are *bona fide* products of splicing. Importantly, these spliced transcripts can be detected associated to high order polysomes, which strongly supports their active translation (Figure 1H).

To assess the relevance of splicing for the viral infection, we inoculated *SF3B1*-silenced plants with TYLCV; as shown in Figure 2A and B, *SF3B1*-knock-down, which compromises viral splicing (Figure 1E, F), limits viral accumulation, pointing at a requirement of splicing for full infectivity. However, the possibility that impaired splicing of host transcripts contributes to this negative effect on infection cannot be excluded.

**Figure 2.**
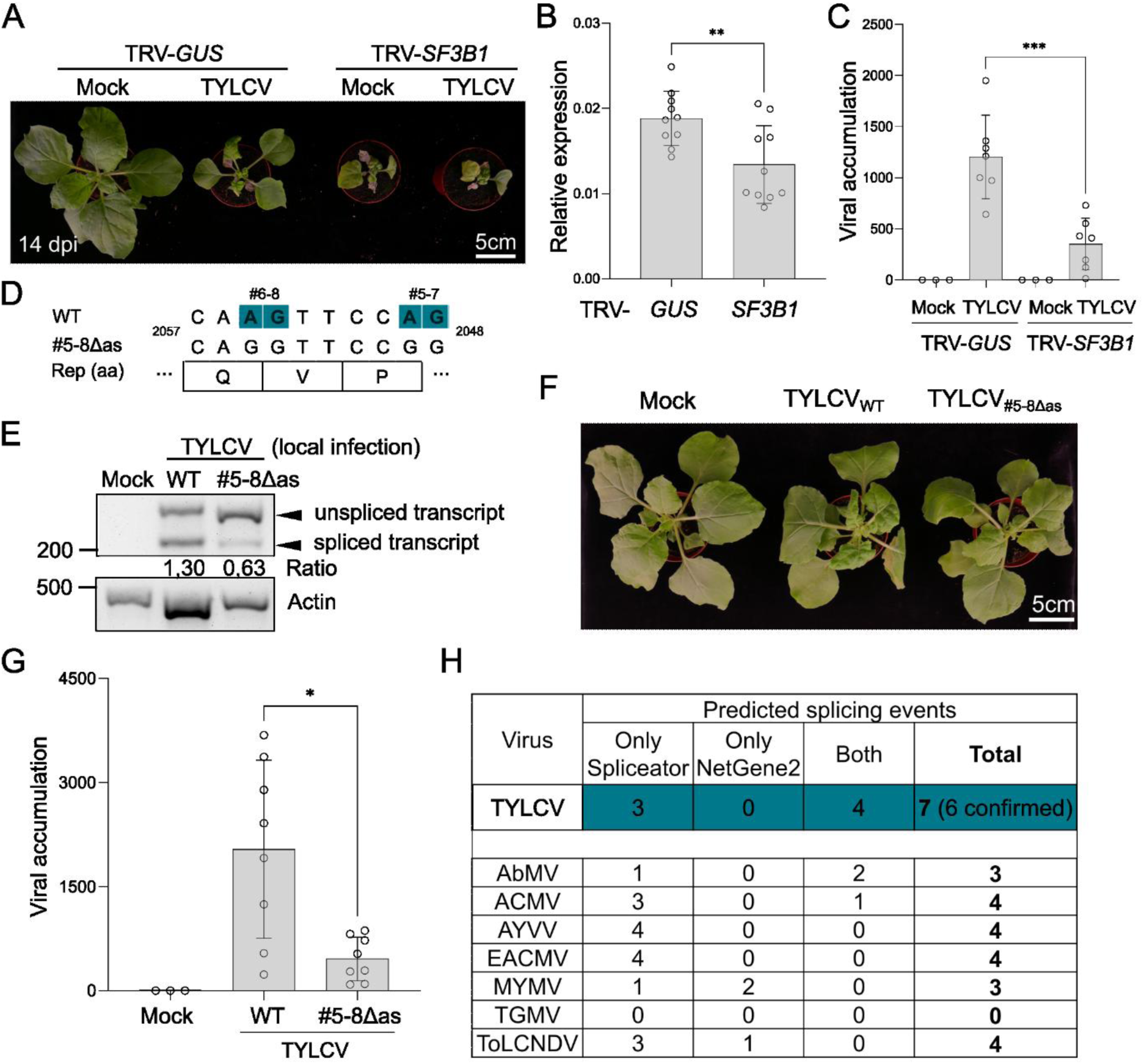
Splicing is required for viral infectivity. (A) Mock- and TYLCV-inoculated TRV- *GUS-* and TRV-*SF3B1-*infected plants (14 days post-inoculation, dpi). (B) *SF3B1* relative expression in the mock- and TYLCV-inoculated plants in A. (C) TYLCV accumulation in the plants in A. (D) Mutations were introduced in the 3’ splice sites for splicing events #5-8 (in blue), without affecting the Rep protein sequence. Position of the alignment (in nt) is shown. (E) RT-PCR showing accumulation of Rep spliced transcript (event #5) in TYLCV_WT_- and TYLCV_#5-8Δas_-inoculated *N. benthamiana* plants (local infection, 3 dpi). The spliced-to-unspliced transcript ratio is shown. (F) Mock-, TYLCV_WT_- and TYLCV_#5-8Δas_-inoculated plants (14 dpi). (G) TYLCV accumulation in the plants in F. (H) Splicing prediction in the Rep/CP transcripts of different begomoviruses, using Spliceator (Scalzitti et al., 2021) and NetGene2 (Hebsgaard et al., 1996). For TYLCV, 6/7 predicted splicing events were experimentally identified (Supplementary table 1). ACMV: African cassava mosaic virus; AbMV: abutilon mosaic virus; AYVV: *Ageratum* yellow vein virus; EACMV: East African cassava mosaic virus; MYMV: mungbean yellow mosaic virus; TGMV: tomato golden mosaic virus: ToLCNDV: tomato leaf curl New Dehli virus. Both: splicing event was identified with Spliceator and NetGene2 predictor tools.

We next focused on splicing event #5 to evaluate the relevance of this spliced variant during infection. We generated a TYLCV mutant in which the acceptor splice sites are modified, introducing a synonymous substitution in the Rep gene, without affecting the coding sequence of other overlapping reading frames (i.e. C4). This mutation is predicted to interfere with splicing events #5-8 on the Rep transcript, which lead to the removal of the central domain of the Rep protein. Hereafter, we refer to this mutant as TYLCV_#5-8Δas_ (for *acceptor sites in splicing events #5-8*; Figure 2D; Figure 1A; Supplementary table 1). This mutation largely reduces the occurrence of splicing event #5 in locally-infected *N. benthamiana* leaf patches (Figure 2E). Interestingly, TYLCV_#5-8Δas_ shows significantly decreased accumulation in systemic infections in *N. benthamiana* plants (Figure 2G), which mostly remain asymptomatic (Figure 2F). We can therefore conclude that splicing is essential for TYLCV pathogenicity.

Our results show that pervasive splicing affects transcripts of the geminivirus TYLCV in the context of the viral infection, indicating that splicing of viral transcripts is much more prevalent than previously thought. Importantly, viral splicing i) affects transcripts encoded both in the viral and in the complementary strands of the viral genome (Figure 1A, B; Supplementary table 1); ii) requires the plant spliceosome (Figure 1C-E); iii) generates transcripts associated to high-order polysomes and is predicted to give rise to new protein isoforms (Figure 1H; Supplementary table 1); iv) does not require virus-encoded proteins, since transient expression of viral genes in isolation is sufficient to produce spliced transcripts (Figure 1E); and v) contributes to full infectivity (Figure 2F, G). Only a small fraction of viral transcripts undergoes splicing (0.42 to 1.62% according to our short-read RNA-seq data; Supplementary table 2), which might have hampered the discovery of these events. Whether other geminivirus species exhibit a similar degree of transcript splicing remains to be determined, but the notion is supported by splicing predictions across the *Begomovirus* genus (Figure 2H; Supplementary Figure 2 for CP and Pott et al. (2026) for Rep transcripts).

Our findings suggest that splicing represents an additional, so far overlooked strategy by which geminiviruses expand their proteome and functional repertoire. The subset of proteins generated by splicing adds to those encoded by novel, previously unrecognized genes recently reported in this viral family (Tan et al., 2025). TYLCV was long believed to encode six proteins (Rep, C2, C3, C4, V2, and CP), but three more (V3, C5, and C7) have been recently discovered (Gong et al., 2021; Liu et al., 2023; Zhao et al., 2022). Our data suggest that splicing could produce at least eight additional protein isoforms, effectively doubling the currently accepted TYLCV proteome.

While our proof of concept demonstrates that viral splicing is essential for full infection, a systematic dissection of individual splicing events will be required to clarify their specific contributions to the viral cycle. The characterization of the resulting protein isoforms will provide new insights into viral infection dynamics and virus–host interactions, and may uncover novel targets for resistance. Considering these findings, we propose that the transcriptomes and proteomes of plant viruses, particularly those with nuclear-replicating DNA genomes, should be revisited. By revealing pervasive splicing in a plant virus, our work propels a paradigm shift in our understanding of the plant-virus interface, uncovering an overlooked dimension which will be key to understanding, and ultimately controlling, plant virus diseases.

## METHODS

### Plant material

*Nicotiana benthamiana* and *Solanum lycopersicum* (cv. “Moneymaker” – viral infections – and “Santa Clara” – ribosome profiling) were grown in a controlled growth chamber under long-day conditions (16 h of light/8 h of dark) at 25 °C.

### Bacterial strains and growth conditions

*Escherichia coli* strain DH5α was used for general cloning and subcloning procedures; strain DB3.1 was used to amplify Gateway-compatible empty vectors. *Agrobacterium tumefaciens* strain GV3101 harboring the corresponding binary vectors was used for *in planta* expression.

### Plasmid construction

All primers used in this study are summarized in Dataset S1. The TYLCV clone used as template is AJ489258 (NCBI:txid220938); the CpCDV clone used as template is DQ458791 (NCBI:txid463360). TYLCV_#5-8Δas_ was generated by site-directed mutagenesis PCR, with the corresponding primers (Supplementary table 3). The DNA fragment carrying the mutations was then cloned into the pDONR/Zeo entry vector (Thermo Scientific) and was further recombined into pGWB501 (Nakagawa, Kurose, et al., 2007; Nakagawa, Suzuki, et al., 2007) destination vector through a Gateway LR reaction (Thermo Scientific).

Open reading frames of *Rep* and *CP* from TYLCV were cloned into pDONR/Zeo entry vector and introduced to pGWB502 by a Gateway LR reaction (Nakagawa, Kurose, et al., 2007; Nakagawa, Suzuki, et al., 2007). For virus-induced gene silencing (VIGS) assay, *SF3B1* homolog was identified in *N. benthamiana* (Niben101Scf06716g01016). A 300-bp conserved gene fragment was PCR-amplified, and cloned into the pTRV2 vector (Liu et al., 2002) by traditional cloning using *Bam*HI and *Kpn*I.

### *Agrobacterium tumefaciens*-mediated transient gene expression in *N. benthamiana*

*A. tumefaciens* cells carrying the constructs of interest were liquid-cultured in LB with the appropriate antibiotics at 28°C overnight. Bacterial cultures were centrifuged at 4,000 *g* for 5 min and resuspended in infiltration buffer (10 mM MgCl_2_, 10 mM MES, pH 5.6, and 150 µM acetosyringone) to an OD_600_ = 0.1–0.5. After a 2h incubation at room temperature in the dark, bacterial cultures were used to infiltrate the abaxial side of leaves of 3- to 4-week-old *N. benthamiana* with a 1 mL needleless syringe. Plants were kept in the greenhouse for 24-48h before further analysis.

### Virus-induced gene silencing (VIGS) assay

VIGS assays were performed as described (Yu et al., 2019). Briefly, independent cultures of *A. tumefaciens* carrying pTRV1 or pTRV2-based constructs were liquid-cultured overnight in LB medium with appropriate antibiotics. Cultures were resuspended in 10 mM MgCl_2_, 10 mM MES pH 5.6 and 150 mM acetosyringone to OD_600_ = 0.5, and incubated for 2 h at room temperature in the dark. Cultures were mixed at a 1:1 ratio. Approximately 1 mL of this suspension was used to inoculate the stem and underside of cotyledons of two-week-old *N. benthamiana*.

### Virus infection assays

For TYLCV and CpCDV local infection assays, fully expanded young leaves of 4-week-old *N*. *benthamiana* plants were infiltrated with *A*. *tumefaciens* carrying TYLCV infectious clones. Samples were collected at 3 days post-inoculation (dpi) to detect viral accumulation and viral gene splicing. For TYLCV systemic infection assays, the stems of 2-week-old *N. benthamiana* or tomato plants were syringe-inoculated with *A*. *tumefaciens* carrying TYLCV infectious clones. The youngest apical leaves were harvested at 14 dpi to detect viral accumulation. For VIGS of *SF3B1*, plants were co-inoculated with a suspension of *A. tumefaciens* carrying TYLCV infectious clone.

### Determination of viral accumulation and quantitative PCR (qPCR)

To determine viral accumulation, total DNA was extracted from *N*. *benthamiana* leaves using the CTAB method (Murray & Thompson, 1980). Quantitative PCR (qPCR) was performed with primers to amplify TYLCV *Rep(Wang et al., 2017)*. As internal reference for DNA detection, the *25S ribosomal DNA interspacer* (*ITS*) was used (Mason et al., 2008). qPCR was performed in a BioRad CFX384 real-time system with PowerTrack SYBR Green Mastermix (Thermo Scientific) with the following program: 2 min at 95°C, and 40 cycles consisting of 15 s at 95°C, 1 min at 60°C.

### RNA extraction and reverse transcription (quantitative) PCR (RT-(q)PCR)

RNA was extracted following the citrate-citric acid RNA isolation method (Oñate-Sánchez & Verdonk, 2021). RNA was treated with DNAse I (Thermo Scientific), and 500 ng of clean RNA was used for cDNA synthesis, with the PrimeScript RT MasterMix (Takara) following manufacturer’s instructions. The qPCR reaction was performed with PowerTrack SYBR Green Mastermix, following the program: 2 min at 95°C, and 40 cycles consisting of 15 s at 95°C, 1 min at 60°C. *Elongation factor-1 alpha* (*NbEF1α*) (Segonzac et al., 2011) was used as reference gene. For splicing analysis, primers were designed spanning the splice sites to favor the amplification of the spliced transcripts, although unspliced transcripts were also amplified. In the case of splicing event #2, only a weak band could be detected for the unspliced transcript, while for splicing event #5 the unspliced transcript is robustly amplified and was used to quantify the spliced to unspliced ratio. Amplified bands were visualized on an agarose gel after PCR, and were quantified with ImageJ software.

### RNA splicing analysis

RNA-seq data of *N. benthamiana* plants infected with TYLCV was used for splicing analysis (Wang et al., 2022). Splicing events were identified using rMATS/rMATS.3.2.2 (Shen et al., 2014) with the following options: -t paired -len 150 -an 8 -c 0.0001 – analysis U -keep temp. Percent Spliced In (PSI) values were calculated for each splicing event. Junction reads are defined as those spanning the splicing site; retention reads are defined as those showing at least a 10-bp overlap with the intron region.

### Polysome profiling

Polysome profiling assay was performed as described(Mustroph et al., 2009), with slight modifications. 1 g of tomato leaves were thawed in Polysome Extraction Buffer (200 mM Tris-HCl pH 9, 200 mM KCl, 200 mM Sucrose, 25 mM EGTA, 35 mM MgCl_2_, 1% detergent mix 1% (20% polyoxythylene(23)lauryl ether (Brij-35), 20% Triton X-100, 20% octyphenyl-polyethylene glycol (Igepal CA 630), 20% Tween 20), 5mM DTT, 50 μL/mL Cycloheximide, 50 μL/mL Chloramphenicol). The extract was cleared by filtering it through three layers of miracloth and by centrifugation at 7000 *g* for 20 min at 4°C. Supernatant was layered onto a sucrose cushion (1,75 M Sucrose, 400 mM Tris-HCl pH 9, 200 mM KCl, 5 mM EGTA, 35 mM MgCl_2_, 5mM DTT 50 μL/mL Cycloheximide, 50 μL/mL Chloramphenicol) and centrifuged for 3,5 h at 40.000 rpm at 4°C in a SW40 rotor. Polysome pellets were resuspended (200 mM Tris-HCl pH 9, 200 mM KCl, 25 mM EGTA, 35 mM MgCl_2_, 5mM DTT, 50 μL/mL Cycloheximide, 50 μL/mL Chloramphenicol) and layered on top of a 12.5 mL ml 10%–50% sucrose density gradient (40 mM Tris-HCl (pH 8.4), 20 mM KCl, 10 mM MgCl_2_, 50 μL/mL Cycloheximide, 50 μL/mL Chloramphenicol) and centrifuged at 36,300 rpm for 2.5 h at 4 °C in a SW40 rotor. After ultracentrifugation, the gradient was monitored at A256 nm while being fractionated into 600 μl fractions using a density gradient fractionation system (BioComp Instruments). RNA from the sucrose fractions was isolated following(Merchante et al., 2016).

### Statistical analysis

Statistical significance was analyzed with GraphPad Prism 9 using unpaired Student’s *t*-test or one-way ANOVA. Significance levels of *p* < 0.05 (*), *p* < 0.01 (**), *p* < 0.001 (***) and *p* < 0.0001 (****) were chosen. Error bars are shown as mean ± standard deviation.

## SUPPLEMENTARY INFORMATION

**Supplementary table 1.** Splicing events detected in TYLCV transcripts in RNA-seq of infected *Nicotiana benthamiana* plants. Intron position, intron size, 5’ and 3’ splice sites, affected viral ORFs, and effects of splicing at the protein level are shown. Protein length of the isoforms is indicated in parentheses.

**Supplementary table 2. Relative quantification of spliced transcripts.** RNA-seq data are from Wang, Tan, et al., 2022, and are deposited in NCBI Gene Expression Omnibus (GEO) under project number GSE309527. Percent Spliced In (PSI) values were calculated for each splicing event. Junction reads are defined as those spanning the splicing site; retention reads are defined as those showing at least a 10-bp overlap with the intron region.

**Supplementary table 3. Primers used in this study.**

**Supplementary figure 1. Alignments of sequenced splicing events #2 and #5 against TYLCV genomic regions 392-763 and 1893-2318, respectively.** Splicing events #2 and #5 from Supplementary table 1 are highlighted in green.

**Supplementary figure 2. Splicing prediction in CP transcripts across the Begomovirus genus.** (A) Predicted splicing events in similar region than splicing event #1 from TYLCV. (B) Predicted splicing events in similar region than splicing event #2 from TYLCV. (C) Predicted splicing events in similar region than splicing event #3 from TYLCV. (D) Additional predicted splicing events. Predictions were performed with Spliceator as described in Pott et al. (2026). AAEO: Africa, Asia, Europe, Oceania; Sweepoviruses: sweet potato-infecting begomoviruses.

**Supplementary figure 3. Prediction and confirmation of a splicing event in the CP transcript from the mastrevirus chickpea chlorotic dwarf virus (CpCDV).** (A) Prediction of the splicing event according to Spliceator. (B) RT-PCR with a reverse primer flanking the intron-exon junction amplifies an 181-bp fragment from cDNA from *N. benthamiana* leaves locally infected with CpCDV at 3 days post-inoculation. Numbers 1 to 7 represent independent biological replicates. Amplified bands from samples 1 and 6 were isolated from the gel and purified for Sanger sequencing to confirm the splicing event. (C) Alignment of the sequenced splicing event with the CP full-length transcript.

## ACKNOWLEDGEMENTS

The authors thank Clémence Marchal, Zhihao Jiang, Axel Giudicatti, Hua Wei, and Huang Tan for critical reading of the manuscript, and Bettina Stadelhofer, Gemma Sans Coll, and the central facilities at the ZMBP, especially the plant cultivation and the microscopy facilities, for excellent technical support. This work was partially funded by the DFG (SFB 1101/C08) and the European Research Council (GemOmics; 101044142); DMP acknowledges previous funding from the Alexander von Humboldt Foundation and is the recipient of a Marie Skłodowska-Curie Grant from the European Union’s Horizon Program (VIRALS; 101104619).

