## Supplementary material for "Pervasive RNA splicing in a plant DNA virus"

[#]Corresponding author

**Supplementary table 1**

**Supplementary table 2**

**Supplementary table 3**

**Supplementary figure 1**

**Supplementary figure 2**

**Supplementary figure 3**

**Supplementary table 1.** Splicing events detected in TYLCV transcripts in RNA-seq of infected *Nicotiana benthamiana* plants. Intron position, intron size, 5’ and 3’ splice sites, affected viral ORFs, and effects of splicing at the protein level are shown. Protein length of the isoforms is indicated in parentheses.

| Splicing Event | Spliced fragment (nt) | Putative intron size (nt) | Spliced isoform (%) | Splice sites | Viral transcripts | Spliced proteins |
|---|---|---|---|---|---|---|
| #1 | 577-669 | 93 | 1.62 | GT-AG | CP, C5 | (227 aa)<br>(110 aa) |
| #2 | 634-745 | 112 | 0.65 | GT-AG | CP, C5 | (132 aa)<br>(44 aa) |
| #3 | 634-748 | 115 | 0.87 | GT-AG | CP, C5 | (131 aa)<br>(43 aa) |
| #4 | 783-876 | 94 | 0.51 | GT-AG | CP, C5 | (163 aa)<br>C5: start codon is spliced out |
| #5 | 2049-2207 | 159 | 0.37 | GC-AG | Rep, C4 | (304 aa)<br>(113 aa) |
| #6 | 2055-2207 | 153 | 0.10 | GC-AG | Rep, C4 | (306 aa)<br>(115 aa) |
| #7 | 2049-2252 | 204 | 1.06 | GT-AG | Rep, C4 | (289 aa)<br>(128 aa) |
| #8 | 2055-2252 | 198 | 0.42 | GT-AG | Rep, C4 | (291 aa)<br>(130 aa) |

Rep -357 aa
CP -258 aa
C4 -97 aa
C5 -67 aa
Frameshift
------ Intron

**Supplementary table 2. Relative quantification of spliced transcripts.** RNA-seq data are from Wang, Tan, et al., 2022, and are deposited in NCBI Gene Expression Omnibus (GEO) under project number GSE309527. Percent Spliced In (PSI) values were calculated for each splicing event. Junction reads are defined as those spanning the splicing site; retention reads are defined as those showing at least a 10-bp overlap with the intron region.

| Splicing event | Junction reads | Retention reads | Total reads | PSI values |
|---|---|---|---|---|
| #1 | 20068 | 1222450 | 1242518 | 0.9838 |
| #2 | 8532 | 1299832 | 1308364 | 0.9935 |
| #3 | 11543 | 1308698 | 1320241 | 0.9913 |
| #4 | 4321 | 838444 | 842765 | 0.9949 |
| #5 | 343 | 92895 | 93238 | 0.9963 |
| #6 | 90 | 92243 | 92333 | 0.9990 |
| #7 | 1306 | 122054 | 123360 | 0.9894 |
| #8 | 511 | 121407 | 121918 | 0.9958 |

**Supplementary table 3. Primers used in this study.** F: forward; R: reverse. In lowercase, attB sites for Gateway compatible cloning. F primer includes attB1 sequence; R primer includes attB2 sequence. Underlined sequences correspond to BamHI and KpnI restriction sites.

| Name | Species | Reference | Primer Sequence (5'-3') | Gene ID |
|---|---|---|---|---|
| **For cloning:** | | | | |
| *Rep* | TYLCV | this study | F: ggggacaagtttgtacaaaaaagcaggctccATGCCTCGTTTATTTAAAAT<br>R: ggggaccactttgtacaagaaagctgggtcTTACGCCTTATTGGTTTCTTCTTG | N/A |
| *CP* | TYLCV | this study | F: ggggacaagtttgtacaaaaaagcaggctccATGTCGAAGCGACCAGGCG<br>R: ggggaccactttgtacaagaaagctgggtcTTAATTTGATATTGAATCATAGAAATAGATGCG | N/A |
| TRV2-*SF3B1* | *Nicotiana benthamiana* | this study | F:CGCGGATCCCAGTATTTTGGTTCCTTGTTGAAC<br>R: CGGGGTACCGTGCACATACGGGCGAA | Niben101Scf06716g01016 |
| **For site directed mutagenesis:** | | | | |
| TYLCV$_{\#5-8\Delta ass}$ | TYLCV | this study | F: CTTCTTCTTTTAATCAGGTTCCGGATGAACTTGAAGAGTGGG<br>R: CCCACTCTTCAAGTTCATCCGGAACCTGATTAAAAGAAGAAG | N/A |
| **For q(RT)-PCR:** | | | | |
| *Rep* | TYLCV | Wang et al., 2017 | F: TGAGAACGTCGTGTCTTCCG<br>R: TGACGTTGTACCACGCATCA | N/A |
| *Internal transcript spacer 25SrDNA (ITS)* | *Nicotiana benthamiana* | Mason et al., 2008 | F: ATAACCGCATCAGGTCTCCA<br>R: CCGAAGTTACGGATCCATTT | N/A |
| *NbEF1α (Elongation factor 1-alpha)* | *Nicotiana benthamiana* | Segonzac et al., 2011 | F: AAGGTCCAGTATGCCTGGGTGCTTGAC<br>R: AAGAATTCACAGGGACAGTTCCAATACCAC | N/A |
| *SF3B1* | *Nicotiana benthamiana* | this study | F: ATGAGACTCCTACGCCTGGT<br>R: TGCACTTCCCATAGTGGCTG | Niben101Scf06716g01016 |
| **For RT-PCR:** | | | | |
| *CP* | TYLCV | Wang et al., 2017 | F: TGGAAGCAGCCCAATGGATT<br>R: GTTCTCGTACTTGGCTGCCT | N/A |
| *splicing event #1* | TYLCV | this study | F: AACAAGCGACGATCATGGAC<br>R:CCTGATTAGTGTGATTCTGCTTC | N/A |

| | | | | |
|---|---|---|---|---|
| *splicing event #2* | TYLCV | this study | F: AATCATTTCCACGCCCGTCT<br>R: CTGCTTCCATAGGGCCTTCCACT | N/A |
| *splicing event #3* | TYLCV | this study | F: CCAGTCTTATGAGCAACGGG<br>R:AAATCCATTGGGCTGCTTCC | N/A |
| *splicing event #4* | TYLCV | this study | F:GCCCAATGGATTTTGGACAG<br>R:GTACTTGGCTGCCTCCTGAT | N/A |
| *splicing event #5* | TYLCV | this study | F: GACCCACTCTTCAAGTTCATCTGAAT<br>R: CAACGGTTCTTCGACCTGGT | N/A |
| *splicing event #5 (Sanger sequencing)* | TYLCV | Wang et al., 2017; Wang et al., 2022 | F: ATCCGAACATTCAGGCAGCT<br>R: TGACGTTGTACCACGCATCA | N/A |
| *splicing event #8* | TYLCV | this study | F: CAACGGTTCTTCGACCTGGT<br>R: CTCTTCAAGTTCATCTGGAA | N/A |
| *CP* | CpCDV | this study | F: GAAGAAAACGTGGTCATGCG<br>R: TCCAACTGACTTGAAGTACACCGTTG | N/A |
| *NbActin* | *Nicotiana benthamiana* | this study | F: ATGTTCACCACCTCAGCTGA<br>R: GGGAAGCCAAGATAGAGC | N/A |

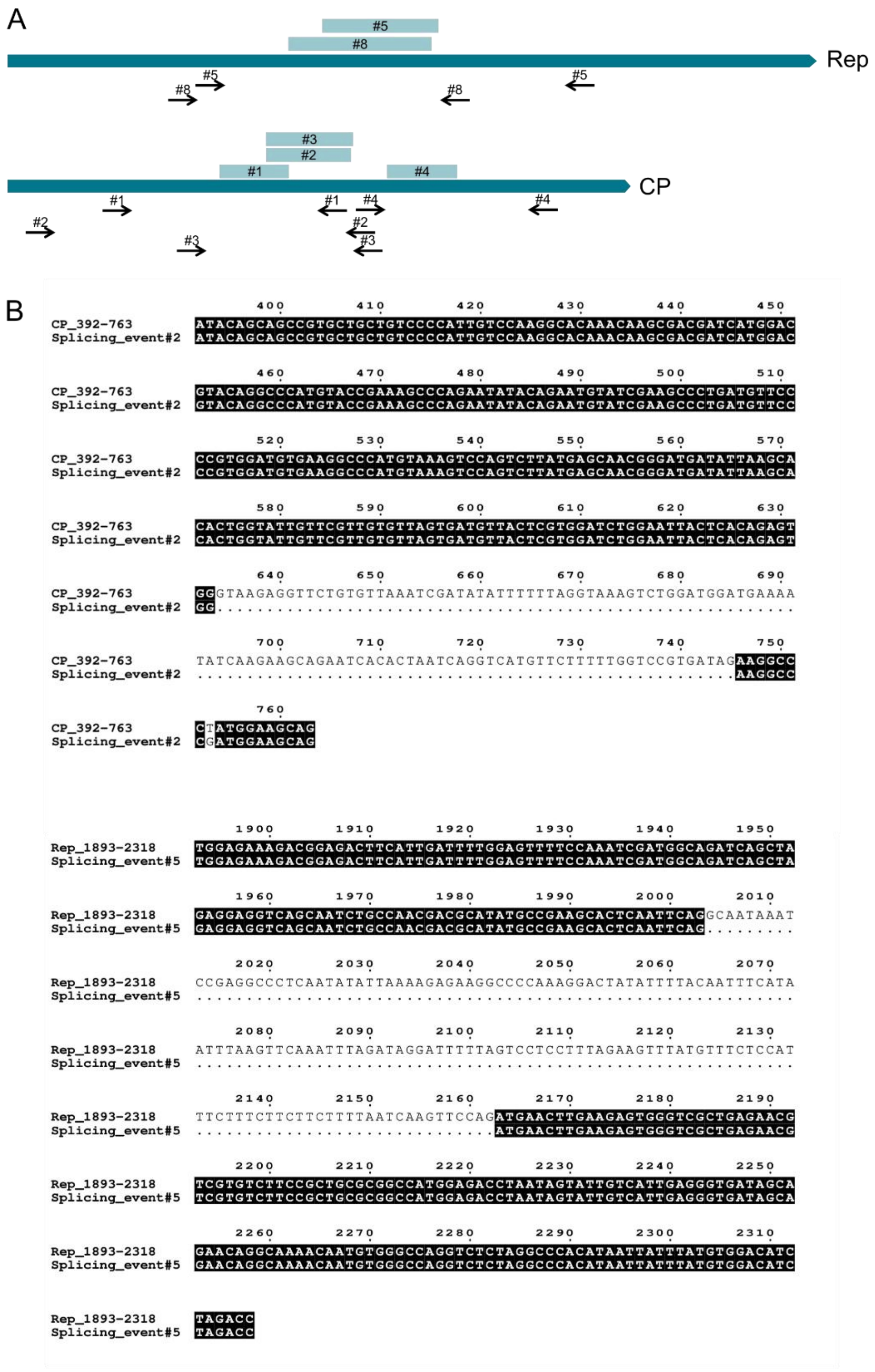
A
#5
#8
Rep
#5
#8
#8
#5
#3
#2
#1
#4
CP
#1
#1
#4
#4
#2
#2
#3
#3
B
400 410 420 430 440 450
CP_392-763 ATACAGCAGCCGTGCTGCTGTCCCCATTGTCCAAGGCACAAACAAGCGACGATCATGGAC
Splicing_event#2 ATACAGCAGCCGTGCTGCTGTCCCCATTGTCCAAGGCACAAACAAGCGACGATCATGGAC
460 470 480 490 500 510
CP_392-763 GTACAGGCCCATGTACCGAAAGCCCAGAATATACAGAATGTATCGAAGCCCTGATGTTCC
Splicing_event#2 GTACAGGCCCATGTACCGAAAGCCCAGAATATACAGAATGTATCGAAGCCCTGATGTTCC
520 530 540 550 560 570
CP_392-763 CCGTGGATGTGAAGGCCCATGTAAAGTCCAGTCTTATGAGCAACGGGATGATATTAAGCA
Splicing_event#2 CCGTGGATGTGAAGGCCCATGTAAAGTCCAGTCTTATGAGCAACGGGATGATATTAAGCA
580 590 600 610 620 630
CP_392-763 CACTGGTATTGTTCGTTGTGTTAGTGATGTTACTCGTGGATCTGGAATTACTCACAGAGT
Splicing_event#2 CACTGGTATTGTTCGTTGTGTTAGTGATGTTACTCGTGGATCTGGAATTACTCACAGAGT
640 650 660 670 680 690
CP_392-763 GGGTAAGAGGTTCTGTGTTAAATCGATATATTTTTTAGGTAAAGTCTGGATGGATGAAAA
Splicing_event#2 GG..........................................................
700 710 720 730 740 750
CP_392-763 TATCAAGAAGCAGAATCACACTAATCAGGTCATGTTCTTTTTGGTCCGTGATAGAAGGCC
Splicing_event#2 ......................................................AAGGCC
760
CP_392-763 CTATGGAAGCAG
Splicing_event#2 CGATGGAAGCAG
1900 1910 1920 1930 1940 1950
Rep_1893-2318 TGGAGAAAGACGGAGACTTCATTGATTTTGGAGTTTTCCAAATCGATGGCAGATCAGCTA
Splicing_event#5 TGGAGAAAGACGGAGACTTCATTGATTTTGGAGTTTTCCAAATCGATGGCAGATCAGCTA
1960 1970 1980 1990 2000 2010
Rep_1893-2318 GAGGAGGTCAGCAATCTGCCAACGACGCATATGCCGAAGCACTCAATTCAGGCAATAAAT
Splicing_event#5 GAGGAGGTCAGCAATCTGCCAACGACGCATATGCCGAAGCACTCAATTCAG.........
2020 2030 2040 2050 2060 2070
Rep_1893-2318 CCGAGGCCCTCAATATATTAAAAGAGAAGGCCCCAAAGGACTATATTTTACAATTTCATA
Splicing_event#5 ............................................................
2080 2090 2100 2110 2120 2130
Rep_1893-2318 ATTTAAGTTCAAATTTAGATAGGATTTTTAGTCCTCCTTTAGAAGTTTATGTTTCTCCAT
Splicing_event#5 ............................................................
2140 2150 2160 2170 2180 2190
Rep_1893-2318 TTCTTTCTTCTTCTTTTAATCAAGTTCCAGATGAACTTGAAGAGTGGGTCGCTGAGAACG
Splicing_event#5 ..............................ATGAACTTGAAGAGTGGGTCGCTGAGAACG
2200 2210 2220 2230 2240 2250
Rep_1893-2318 TCGTGTCTTCCGCTGCGCGGCCATGGAGACCTAATAGTATTGTCATTGAGGGTGATAGCA
Splicing_event#5 TCGTGTCTTCCGCTGCGCGGCCATGGAGACCTAATAGTATTGTCATTGAGGGTGATAGCA
2260 2270 2280 2290 2300 2310
Rep_1893-2318 GAACAGGCAAAACAATGTGGGCCAGGTCTCTAGGCCCACATAATTATTTATGTGGACATC
Splicing_event#5 GAACAGGCAAAACAATGTGGGCCAGGTCTCTAGGCCCACATAATTATTTATGTGGACATC
Rep_1893-2318 TAGACC
Splicing_event#5 TAGACC

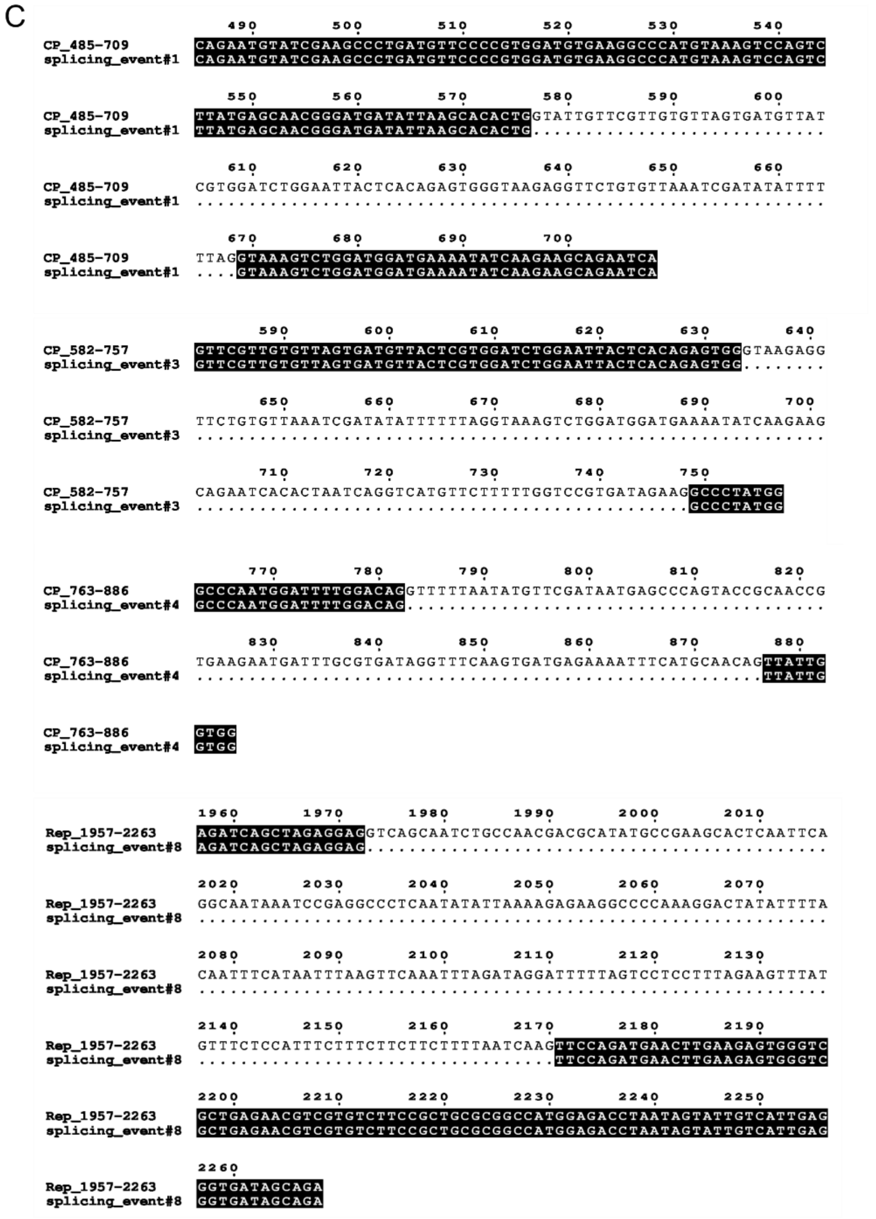

**Supplementary figure 1.** (A) Schematic showing Rep and CP genes with the corresponding splicing events (#1-4 for CP and #5 and #8 for Rep) and primers used for Sanger sequencing and confirmation of the splicing events (black arrow). (B) Alignments of sequenced splicing events #2 and #5 against TYLCV genomic regions. (C) Alignments of sequenced splicing events #1, #3, #4 and #8 against TYLCV genomic regions.

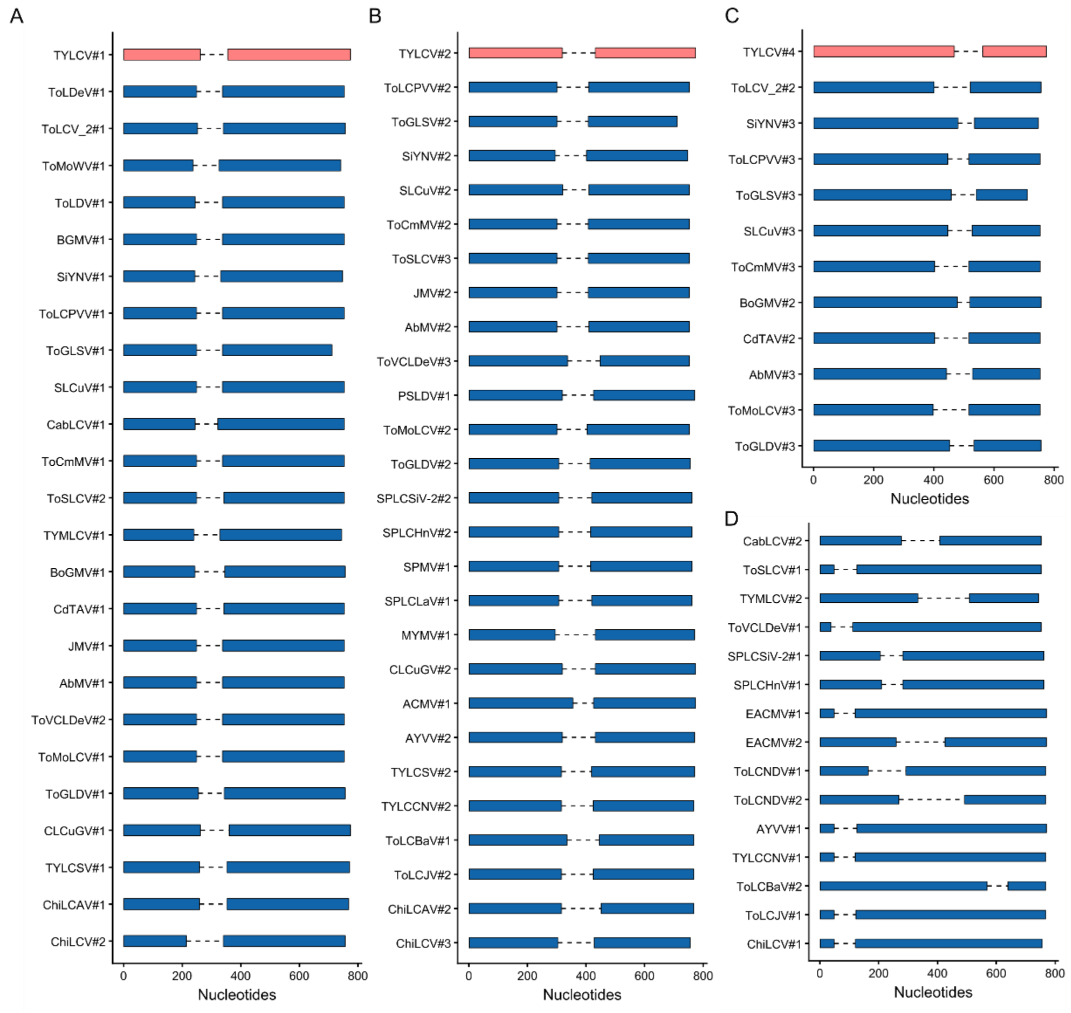


**Supplementary figure 2. Splicing prediction in CP transcripts across the Begomovirus genus.** (A) Predicted splicing events in similar region than splicing event #1 from TYLCV. (B) Predicted splicing events in similar region than splicing event #2 from TYLCV. (C) Predicted splicing events in similar region than splicing event #3 from TYLCV. (D) Additional predicted splicing events. Predictions were performed with Spliceator as described in Pott et al. (2026). AAEO: Africa, Asia, Europe, Oceania; Sweepoviruses: sweet potato-infecting begomoviruses.

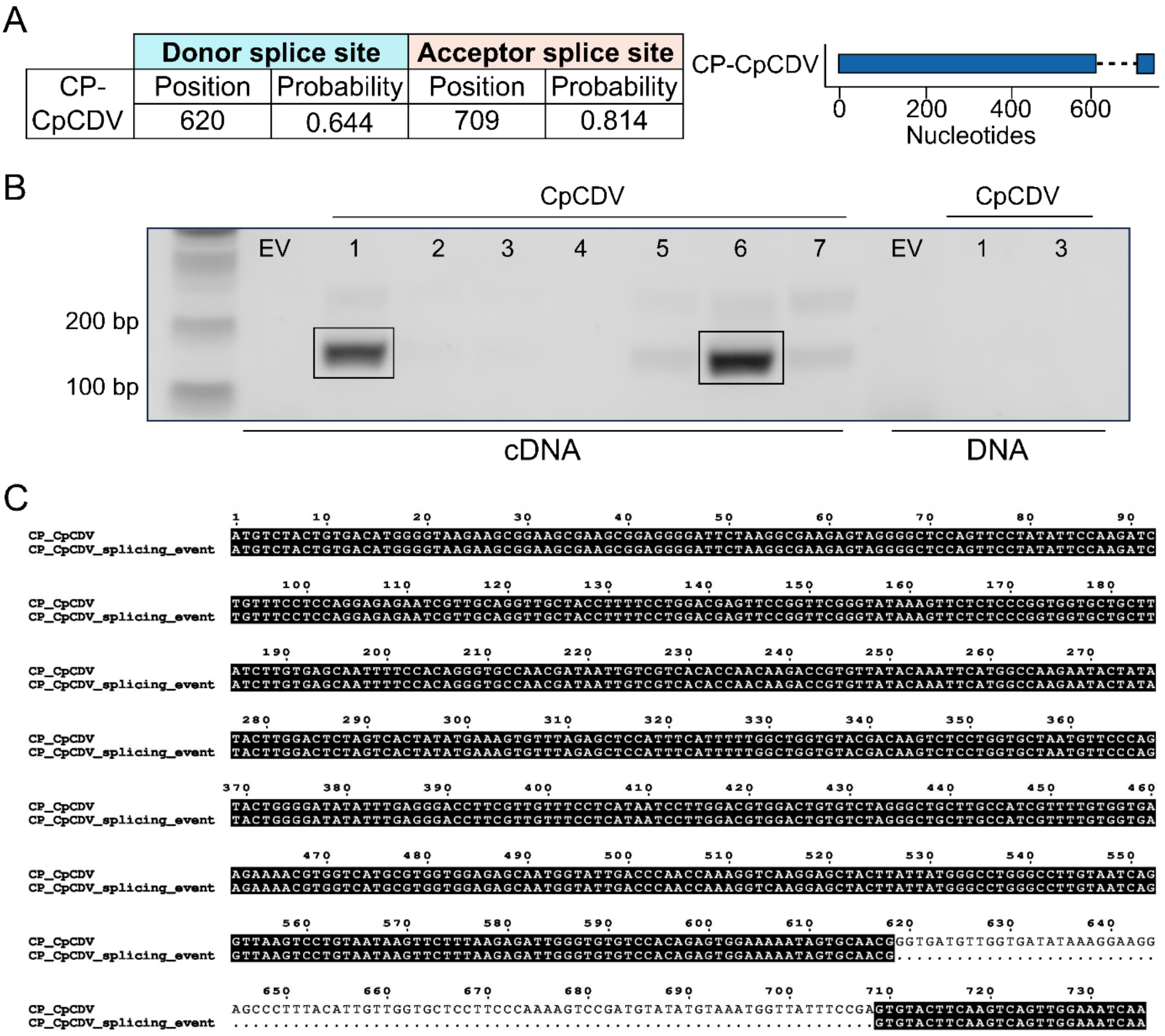


**Supplementary figure 3. Prediction and confirmation of a splicing event in the CP transcript from the mastrevirus chickpea chlorotic dwarf virus (CpCDV).** (A) Prediction of the splicing event according to Spliceator. (B) RT-PCR with a reverse primer flanking the intron-exon junction amplifies an 181-bp fragment from cDNA from *N. benthamiana* leaves locally infected with CpCDV at 3 days post-inoculation. Numbers 1 to 7 represent independent biological replicates. Amplified bands from samples 1 and 6 were isolated from the gel and purified for Sanger sequencing to confirm the splicing event. (C) Alignment of the sequenced splicing event with the CP full-length transcript.
